# Pan-cancer single-cell analysis identifies a FOXF1/FOXF2-associated transitional CAF-like fibroblast state

**DOI:** 10.64898/2026.07.09.737397

**Authors:** Bayrta Mandzhieva, Ayushi Verma, Truc Do Thanh Nguyen, Yeong Hak Bang, Woong-Yang Park

## Abstract

Fibroblast heterogeneity shapes tumor progression, yet the transitional states linking normal-associated fibroblasts to cancer-associated fibroblasts (CAFs) remain poorly defined. Here, we integrated single-cell transcriptomic profiles of more than 90,000 stromal cells from 281 samples across nine cancer types to construct a pan-cancer atlas of fibroblast diversity. We identified a distinct CAF-like population positioned between normal-activated fibroblasts and established CAF subsets along the inferred fibroblast activation trajectory. Integration with single-nucleus chromatin accessibility data nominated FOXF1 and FOXF2 as candidate regulators of this CAF-like state. Functionally, CAF-like fibroblasts were characterized by non-canonical WNT activity, WNT5A-associated stromal communication, and a candidate GZMA–F2R/PAR immune–stromal signaling axis supported by spatial transcriptomics. Clinically, the CAF-like signature showed context-dependent prognostic relevance, with high expression associated with poorer survival in the tumor compartment of TCGA stomach adenocarcinoma. Together, this study defines a FOXF1/FOXF2-associated CAF-like transitional fibroblast state and links it to stromal signaling, immune–stromal communication, and cancer-type-specific clinical relevance.

## Introduction

The tumor microenvironment (TME) is a complex and dynamic ecosystem composed of malignant cells, immune populations, endothelial cells, pericytes, smooth muscle cells, fibroblasts, and extracellular matrix components^1,2^. These non-malignant stromal elements are not passive bystanders; instead, they actively influence tumor growth, invasion, immune regulation, metastatic dissemination, and therapeutic response^3,4,5^. Among these stromal populations, cancer-associated fibroblasts (CAFs) represent one of the most abundant and functionally diverse cell types within solid tumors^6,7^. Through secretion of extracellular matrix proteins, growth factors, cytokines, chemokines, and remodeling enzymes, CAFs shape the physical and signaling architecture of the tumor niche and contribute to multiple aspects of cancer progression^8–12^. Despite their importance, CAFs remain difficult to define as a single functional entity. Multiple CAF subtypes have been described across cancer types, including myofibroblastic, inflammatory, antigen-presenting, vascular, matrix-associated, cycling, and developmental CAF states^13–16^. These subtypes differ in marker expression, spatial localization, regulatory programs, and inferred biological roles. This heterogeneity has important clinical implications because CAFs can exert both tumor-promoting and tumor-restraining functions. While many CAF programs support invasion, immune modulation, extracellular matrix remodeling, and therapy resistance, experimental depletion of specific CAF populations has, in some settings, accelerated tumor progression and worsened survival^17–19^. Similarly, several studies have suggested that certain fibroblast subsets may restrain tumor growth or maintain tissue organization^17,20,21^. These observations indicate that CAF biology is highly context dependent and that broad classification of fibroblasts as uniformly tumor-promoting is likely insufficient. The complexity of CAF biology is further shaped by cellular origin, tissue context, cancer type, tumor stage, and local microenvironmental cues^22,23^. CAFs may arise from tissue-resident fibroblasts, pericytes, mesenchymal progenitors, endothelial-to-mesenchymal transition, bone marrow-derived cells, or other stromal sources. However, whether CAFs emerge as distinct pre-existing lineages or through progressive activation of normal-associated fibroblasts remains incompletely understood. This question is particularly important because most studies focus on established CAF states within tumor tissue, whereas fibroblasts in adjacent non-malignant compartments are often less deeply resolved. As a result, the intermediate states that may connect normal-associated fibroblasts to mature CAF phenotypes remain poorly characterized.

Single-cell RNA sequencing has transformed the study of fibroblast heterogeneity by enabling the identification of distinct stromal cell states at high resolution^24,25^. Recent single-cell studies have revealed multiple CAF subtypes across individual tumor types and have highlighted substantial variability in CAF programs across malignancies^26,27^. However, many analyses are limited by tumor-type-specific datasets, incomplete integration of adjacent normal tissues, or limited consideration of transitional fibroblast states. A pan-cancer framework that jointly analyzes tumor, adjacent normal, and healthy tissue-derived stromal populations could help clarify whether conserved fibroblast activation states exist across cancers and whether intermediate states contribute to CAF evolution. Another important challenge is to understand the regulatory mechanisms that define fibroblast state identity. Transcription factors and chromatin accessibility programs can stabilize fibroblast phenotypes and influence transitions between normal-associated and tumor-associated states. Prior studies have implicated regulatory factors such as RUNX1, AP-1 family members, NF-κB-related factors, STAT/IRF programs, and KLF family transcription factors in shaping distinct fibroblast identities and activation states^28–32^. However, the regulatory architecture of potential intermediate fibroblast states remains less clear. Whether specific transcriptional and epigenetic programs mark a transitional phase between NAFs and mature CAFs has not been fully resolved. In this study, we integrated single-cell transcriptomic data from stromal cells across multiple cancer types, together with adjacent normal and healthy tissue-derived fibroblast populations, to construct a pan-cancer atlas of fibroblast heterogeneity. We aimed to test the hypothesis that the transition from normal-associated fibroblasts to established CAF states involves a distinct intermediate phase rather than a direct binary switch. Our analysis identified a CAF-like fibroblast population positioned between NAFs and mature CAF subsets along the inferred fibroblast activation trajectory. This CAF-like state was detected across tumor and adjacent normal compartments, suggesting that stromal activation may involve a poised transitional program. To characterize this population, we combined transcriptomic analysis, pseudotime trajectory modeling, transcription factor regulon inference, single-nucleus chromatin accessibility profiling, spatial transcriptomics, cell–cell communication analysis, and TCGA-based survival assessment. We nominate FOXF1 and FOXF2 as candidate regulators associated with the CAF-like state and show that this population is linked to non-canonical WNT activity, WNT5A-associated stromal communication, and a candidate GZMA–F2R/PAR immune–stromal axis. Finally, we show that the prognostic relevance of the CAF-like signature is cancer-type and tissue-context dependent, with a stronger adverse association in the tumor compartment of stomach adenocarcinoma. Together, our findings provide a framework for understanding CAF evolution as a dynamic, context-dependent process and highlight candidate regulatory and communication programs for future stromal-focused investigation.

## Materials and Methods

### scRNA-seq data processing

scRNA-seq datasets were obtained from publicly available databases, spanning nine cancer types: colorectal cancer (CRC), lung adenocarcinoma (LUAD), non-small cell lung cancer (NSCLC), hepatocellular carcinoma (LIHC), intrahepatic cholangiocarcinoma (CHOL), pancreatic ductal adenocarcinoma (PDAC), stomach adenocarcinoma (STAD), esophageal squamous cell carcinoma (ESCC), and ovarian cancer (OVC), as well as healthy donor-derived lung, pancreatic, and liver tissues (Supplementary Table 1). Stromal cells were identified in each dataset based on the expression of canonical lineage markers, including DCN and COL3A1 for fibroblasts, PECAM1 and VWF for endothelial cells, ACTG2 and DES for smooth muscle cells, and RGS5 and MCAM for pericytes. Stromal cells from all datasets were then integrated into a single dataset for downstream analyses.

Gene expression matrices were analyzed using the Seurat R package (version 5.0.1)^33^. Quality control filtering was applied, retaining cells with a mitochondrial gene percentage ≤ 20% and a detected gene count between 200 and 6,000. Gene expression data were log-normalized using the NormalizeData function with a scale factor of 10,000. The top 3,000 highly variable genes were identified using FindVariableFeatures (VST selection method), and their expression values were scaled using ScaleData. Principal component analysis (PCA) was conducted on the variable features, retaining the top 100 principal components. To account for batch effects across datasets, the Harmony algorithm was applied to the PCA embedding with tumor type as the grouping variable^34^. Dimensionality reduction was subsequently performed using Uniform Manifold Approximation and Projection (UMAP) on the first 40 Harmony embeddings. Graph-based clustering was conducted by constructing a shared nearest neighbor (SNN) graph using FindNeighbors, followed by modularity optimization using FindClusters at a resolution of 0.5. Stromal cell populations were annotated using the marker genes described above. Integration quality was quantitatively assessed by Local Inverse Simpson’s Index (LISI), calculated with tumor type as the grouping variable before and after Harmony correction. Higher LISI scores indicate improved mixing of cells across datasets.

To investigate fibroblast heterogeneity, the fibroblast population was extracted and re-clustered using the same pipeline described above. This analysis identified eight transcriptionally distinct fibroblast subpopulations, which were annotated based on the expression of subtype-specific marker genes: myofibroblast-like CAFs (myCAF), characterized by TAGLN, ACTA2, MMP11, COL1A2, and COL10A1; inflammatory CAFs (iCAF), marked by CXCL1, CXCL8, IL24, MMP1, and MMP3; interferon-response CAFs (ifnCAF), defined by CXCL10, CXCL11, IFIT2, IFIT3, and ISG15; three normal-activated fibroblast subsets (NAF1, marked by C7, CFD, and IGF1; NAF2, marked by PI16 and PCOLCE2; NAF3, marked by FOS and JUN); normal fibroblasts (NF); and a CAF-like population marked by APOE, A2M, CXCL14, and BMP4.

### Identification of differentially expressed genes

Differential expression analysis was performed using the FindAllMarkers function in Seurat to identify marker genes for each fibroblast subpopulation. Only positively enriched markers were retained (only.pos = TRUE). Genes were required to be expressed in at least 25% of cells within the target cluster (min.pct = 0.25) and to exhibit a minimum log_2_ fold change of 0.25 (logfc.threshold = 0.25). Statistical significance was assessed using the Wilcoxon rank-sum test, and P values were adjusted for multiple comparisons using the Bonferroni correction. Genes with an adjusted P value < 0.05 were considered significantly enriched markers for each fibroblast subpopulation and were used for all downstream analyses.

### Tissue enrichment analysis

To assess the tissue-specific and cancer-type-specific enrichment of fibroblast subpopulations, the ratio of observed to expected cell counts (Ro/e) was calculated for each subtype, as previously described by Zhang et al.^35^. Enrichment was evaluated across two dimensions: tissue compartment (tumor, adjacent normal, and healthy donor) and individual cancer type. For each combination of fibroblast subtype and tissue or cancer category, the expected cell count was calculated as (row total × column total)/grand total, and the Ro/e value was defined as the observed cell count divided by the expected count, with values greater than 1 indicating enrichment.

### Functional analysis

To characterize biological processes associated with each fibroblast subpopulation, Gene Ontology (GO)^36^ analysis was performed using the clusterProfiler R package with the org.Hs.eg.db gene annotation database. For each fibroblast subpopulation, the top 100 DEGs ranked by average log_2_ fold change and filtered at an adjusted P value < 0.05 were converted from gene symbols to Entrez IDs and subjected to GO analysis using the compareCluster function with enrichGO. GO enrichment results were visualized using the dotplot function.

To compare pathway activity between CAF and NAF populations, Gene Set Variation Analysis (GSVA)^37^ was performed using the GSVA R package with Hallmark gene sets retrieved from the Molecular Signatures Database (MSigDB) via the msigdbr R package (Homo sapiens, category “H”). Log-normalized expression data from CAF and NAF cells were used to calculate GSVA enrichment scores using the default Gaussian kernel method. Differential pathway activity was assessed using the limma R package with false discovery rate (FDR) correction, and pathways were ranked according to the t statistic.

To assess pathway-level enrichment specific to the CAF-like population, Gene Set Enrichment Analysis (GSEA)^38^ was performed using the fgsea R package with the same Hallmark gene sets. Gene-level scores for the CAF-like population were computed using the wilcoxauc function from the presto R package and ranked by area under the curve (AUC). Enrichment analysis was then performed using the fgseaMultilevel function. Pathways with an adjusted P value < 0.05 were considered statistically significant. Both analyses were visualized as horizontal bar plots using ggplot2.

### Bulk transcriptomic assessment

To validate fibroblast subpopulation-associated expression patterns identified by scRNA-seq, raw RNA-sequencing count data from the TCGA-COAD cohort were retrieved using the TCGAbiolinks R package with the STAR-Counts workflow. Only primary tumor and solid tissue normal samples were retained for downstream analysis. Raw counts were normalized to log_2_(CPM + 1) using the cpm function from the edgeR R package with a prior count of 1. Selected DEGs representing the CAF, NAF, and CAF-like populations were extracted from the normalized expression matrix, and their expression distributions were compared between tumor and normal samples. Density plots were generated using ggplot2.

### Gene regulatory network inference using SCENIC

To infer transcriptional regulatory networks and transcription factor (TF) activity, SCENIC^39^ (version 1.3.1) was applied to the fibroblast scRNA-seq dataset. To reduce computational complexity, SCENIC analysis was performed on a randomly sampled subset of 10,000 fibroblast cells while preserving the relative abundance of fibroblast subpopulations. The workflow consisted of three main steps: inference of TF–target co-expression modules using GENIE3, cis-regulatory motif enrichment analysis using RcisTarget to identify TF-target regulons, and quantification of regulon activity using the AUCell algorithm. The RcisTarget human motif database with v9 motif annotations (motifAnnotations_hgnc_v9) was used for motif enrichment analysis. Regulon activity scores were calculated for individual cells based on the area under the recovery curve, and regulon activity matrices were binarized using automatically determined AUCell thresholds to identify active and inactive regulons. Regulon activity profiles were subsequently used to compare transcriptional regulatory states among fibroblast subpopulations.

### Single-nucleus ATAC-seq analysis

Single-nucleus ATAC-seq (snATAC-seq) data from colorectal cancer tissue were processed following the standard ArchR pipeline (version 1.0.3)^40^. Arrow files were generated from fragment files, and quality control filtering was performed by retaining nuclei with a TSS enrichment score ≥ 4. Dimensionality reduction was performed using iterative latent semantic indexing (LSI), followed by graph-based clustering and UMAP embedding. Fibroblast nuclei were identified based on cell-type annotations transferred through integration with the scRNA-seq fibroblast reference dataset using ArchR-based gene integration analysis. The resulting fibroblast ArchR object comprised 7,958 nuclei with a median TSS enrichment score of 8.18. Peak calling was performed in a pseudo-bulk manner by aggregating nuclei within each cluster. Differential chromatin accessibility between fibroblast subpopulations was assessed using the getMarkerFeatures function with a bias-corrected Wilcoxon rank-sum test, accounting for TSS enrichment and log-transformed fragment counts. Transcription factor motif enrichment within subpopulation-specific accessible peaks was evaluated using the peakAnnoEnrichment function, with significant enrichment defined as FDR ≤ 0.01 and log2FC ≥ 2. Transcription factor footprinting analysis was performed to infer TF occupancy patterns at predicted motif sites across fibroblast subpopulations.

### Spatial transcriptomics data analysis

Processed 10x Genomics Visium spatial transcriptomics datasets from colorectal and lung cancer tissues were obtained from the 10x Genomics website. Spatial transcriptomic data were loaded into Seurat using the Load10X_Spatial function and normalized using SCTransform normalization. To estimate the spatial distribution of fibroblast subpopulations, cell-type deconvolution was performed using Robust Cell Type Decomposition (RCTD)^41^ implemented in the spacexr package. Spatial coordinates and expression matrices were extracted from Visium objects and used to construct SpatialRNA objects, with the annotated scRNA-seq fibroblast Seurat object used as the reference dataset. RCTD was performed in doublet mode, and predicted cell-type proportions were incorporated into the spatial metadata. To assess the spatial localization of CellChat-predicted ligand–receptor interactions, the expression patterns of selected ligand–receptor pairs were visualized across tissue sections to examine their distribution within the tumor microenvironment.

### Trajectory Analysis

To investigate fibroblast differentiation trajectories, pseudotime analysis was performed using Monocle2^42^ (version 2.26.0), Monocle3^43^ (version 1.3.1), and Slingshot^44^ (version 2.6.0). Gene expression matrices and cell annotations generated in Seurat were used as input for trajectory inference. For Monocle2 analysis, expression data were converted into a CellDataSet object, followed by dimensionality reduction using the reduceDimension function and cell ordering using the orderCells function. Monocle3 and Slingshot were used to independently infer lineage relationships, and consensus trajectories were defined based on concordant lineage relationships identified across all three approaches. Normal fibroblasts (NF), representing unactivated resident fibroblasts, were designated as the trajectory root. Trajectory structures, cell states, and pseudotime ordering were visualized using the default plotting functions implemented in each respective package.

### Pathway activity analysis using PROGENy

Signaling pathway activity was inferred at single-cell resolution using the PROGENy (Pathway RespOnsive GENes) R package^45^. PROGENy estimates pathway activity from the expression of downstream target genes; scores were computed for 14 canonical signaling pathways using the top 500 pathway-responsive genes. Pathway activity scores were subsequently scaled using Seurat’s ScaleData function and averaged across cells within each fibroblast subpopulation. The resulting mean activity scores were visualized as a heatmap using the pheatmap R package with a diverging color scale.

### Module score

To quantify the activity of canonical and non-canonical Wnt signaling programs at single-cell resolution, module scores were computed using the AddModuleScore function in Seurat. Canonical Wnt ligands were defined as WNT2, WNT3, WNT3A, and WNT8A, and non-canonical Wnt ligands as WNT4, WNT5A, WNT5B, WNT6, WNT7A, and WNT11, based on the classification proposed by Qin et al.^46^. Scores were computed as the average expression of each gene set corrected by expression-matched control gene sets and compared across fibroblast subpopulations.

### Cell-cell communication analysis

Intercellular communication networks were inferred using the CellChat R package^47^ (version 1.6.1) separately for the CRC and LC datasets from the pan-cancer scRNA-seq cohort. CellChat objects were constructed from log-normalized expression matrices using the human CellChatDB ligand–receptor database restricted to secreted signaling interactions. Overexpressed genes and ligand–receptor interactions were identified using the identifyOverExpressedGenes and identifyOverExpressedInteractions functions. Gene expression data were projected onto the human protein–protein interaction (PPI) network before communication probability inference. Ligand–receptor interaction probabilities were calculated using computeCommunProb, and pathway-level communication networks were generated using computeCommunProbPathway and aggregateNet. Incoming and outgoing signaling activities were quantified and visualized using heatmaps, and dominant communication patterns were summarized using network-based visualization.

### Survival analysis of a CAF-like gene signature across multiple TCGA cohorts

To evaluate the prognostic relevance of the CAF-like fibroblast population, CAF-like marker genes identified from the scRNA-seq analysis were used to construct a gene signature. A panel of 10 signature genes was selected from CAF-like DEGs ranked by fold change, including APOE, WFDC1, CXCL14, ADAMDEC1, DLL1, LINC01082, HAPLN1, EDNRB, FENDRR, and F2R. RNA-sequencing count data and corresponding clinical information from seven TCGA cohorts (LUAD, COAD, STAD, OV, PAAD, LIHC, and ESCA) were obtained using the TCGAbiolinks R package. Expression data were normalized using the edgeR R package and transformed to log2 counts per million (log2CPM) values. CAF-like signature scores were calculated as the mean log2CPM expression of signature genes detected in each sample. Patients were stratified into high-and low-score groups according to the cohort-specific median signature score.

Overall survival analysis was performed using Kaplan–Meier survival curves and log-rank tests implemented in the survival and survminer R packages. Univariable Cox proportional hazards regression was performed to assess the prognostic association of individual CAF-like signature genes. For the TCGA-STAD cohort, additional analyses were performed separately in tumor and adjacent normal samples to evaluate the tissue-specific prognostic relevance of the CAF-like signature.

## Results

### scRNA-seq analysis reveals fibroblast heterogeneity in tumors

We systematically collected and analyzed single-cell RNA sequencing (scRNA-seq) datasets of stromal cells from nine cancer types (Fig. 1a; Supplementary Table 1). For consistency across related tumor entities, lung adenocarcinoma (LUAD) and non-small cell lung cancer (NSCLC) datasets were grouped as lung cancer (LC), whereas hepatocellular carcinoma (LIHC) and intrahepatic cholangiocarcinoma (CHOL) datasets were grouped as LIHC-CHOL. Healthy donor datasets from lung, pancreas, and liver were also included as non-tumor references to help distinguish normal stromal cell states. Together, the analyzed cohorts covered tumor, adjacent normal, and healthy donor tissues across diverse disease stages and histopathological contexts. After dataset merging and quality-control filtering, the final stromal atlas comprised 90,060 high-quality stromal cells from 281 samples and 234 patients (Fig. 1b). Because the datasets were derived from multiple cancer types and sources, Harmony integration was applied before visualization and clustering to reduce dataset-associated batch effects (Fig. 1c; Supplementary Fig. 1a). Integration quality was assessed using the Local Inverse Simpson’s Index (LISI), with tumor type used as the grouping variable. Higher LISI values after Harmony correction indicated improved local mixing of cells from different tumor types, supporting reduced technical variation across datasets (Supplementary Fig. 1b). Unsupervised graph-based clustering of the integrated stromal cells identified four major stromal populations, which were annotated using canonical marker genes: fibroblasts marked by DCN and COL3A1, endothelial cells marked by PECAM1 and VWF, smooth muscle cells marked by ACTG2 and DES, and pericytes marked by RGS5 and MCAM (Fig. 1d, e). These stromal lineages were broadly represented across cancer types, although their relative proportions varied by tissue context. Fibroblasts formed a major stromal population in most cohorts, whereas endothelial cells were most abundant in the LIHC-CHOL cohort (Fig. 1f), consistent with the highly vascularized architecture of liver tissue and the role of endothelial cells in angiogenic and vascular processes^48^. Given the abundance of fibroblasts and their established roles in tumor progression, we extracted fibroblasts and reclustered them for higher-resolution analysis (n = 50,000). This analysis resolved eight fibroblast subpopulations with distinct transcriptional features (Fig. 1g). These included three cancer-associated fibroblast subsets, -myofibroblast-like CAFs (myCAF), inflammatory CAFs (iCAF), and interferon-response CAFs (ifnCAF); three normal-activated fibroblast subsets (NAF1–3); normal fibroblasts (NF); and a CAF-like population positioned between the CAF and NAF regions in the UMAP embedding. The distribution of these fibroblast states was closely associated with tissue compartment. CAF subsets were preferentially represented in tumor tissues, whereas NAF subsets were enriched in adjacent normal tissues. NF cells were mainly detected in healthy donor samples (Fig. 1h, i). In contrast, CAF-like fibroblasts were detected in both tumor and adjacent normal compartments (Supplementary Fig. 1c, d), suggesting that this population is not restricted to tumor tissue alone but may represent a fibroblast state with mixed compartmental features. To examine whether these single-cell-derived fibroblast signatures were also detectable in patient-level bulk transcriptomic data, we evaluated CAF, NAF, and CAF-like signature scores in the TCGA-COAD cohort. Consistent with the single-cell analysis, the CAF signature was enriched in tumor samples, whereas the NAF signature was higher in adjacent normal samples. The CAF-like signature was detectable in both compartments (Supplementary Fig. 1e), supporting the presence of a CAF-like stromal transcriptional program in both tumor and adjacent normal tissues. We next examined the marker genes and enriched pathways that distinguished each fibroblast subtype (Fig. 1j, k; Supplementary Table 2). myCAFs showed high expression of myofibroblast and extracellular matrix-associated genes, including TAGLN, ACTA2, MMP11, MMP14, COL1A1, COL1A2, and COL10A1, consistent with enrichment of extracellular matrix organization and collagen biosynthesis pathways^49^. iCAFs were characterized by inflammatory chemokines and cytokines, including CXCL1, CXCL8, IL24, and IL11, together with matrix-remodeling genes such as MMP1 and MMP3, supporting an immunoregulatory fibroblast phenotype^49^. The ifnCAF subset showed an interferon-response program marked by CXCL10, CXCL11, IFIT2, IFIT3, and ISG15^27^. In contrast, NAF populations showed features related to tissue maintenance, complement activation, wound repair, peptidase regulation, and stress responses, with representative genes including C7, CFD, IGF1, PI16, PCOLCE2, FOS, and JUN. The CAF-like subset was marked by APOE, A2M, CXCL14, and BMP4, suggesting a distinct stromal program involving protease regulation, stromal communication, and mesenchymal remodeling. Together, these results define a pan-cancer stromal atlas and show that fibroblast states vary markedly across tumor, adjacent normal, and healthy tissues. The detection of a CAF-like population in both tumor and adjacent normal compartments raises the possibility that this state represents an intermediate fibroblast program, a hypothesis explored in the following analyses.

**Figure 1.**
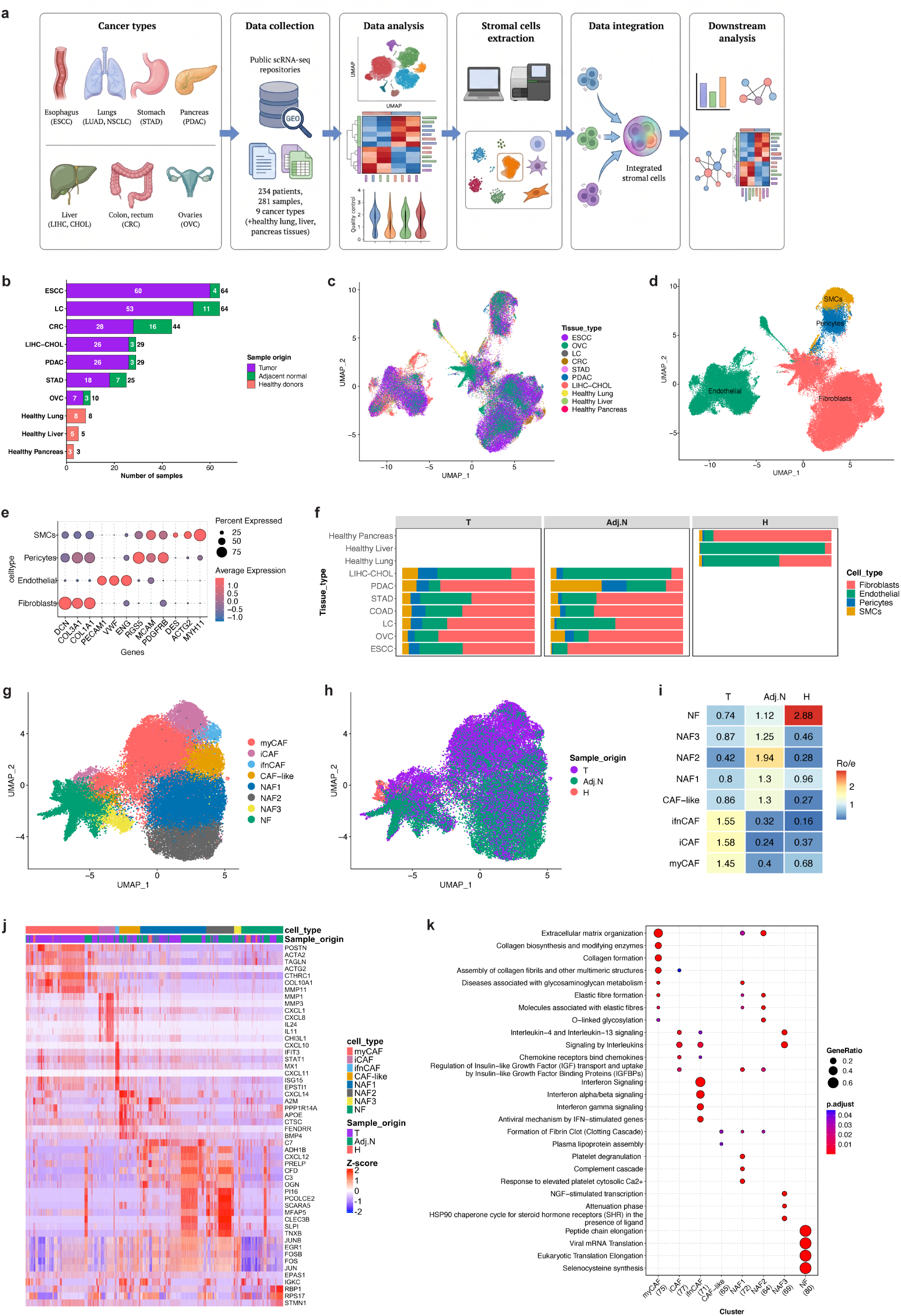
Single-cell landscape of the pan-cancer stromal compartment. **a,** Overview of the workflow used to collect, integrate, and analyze pan-cancer single-cell transcriptomic datasets of stromal cells. **b,** Number of samples included for each cancer or tissue type, stratified by sample origin. **c,** UMAP projection of stromal cells after Harmony integration, colored by cancer or tissue type. **d,** UMAP projection showing the four major stromal lineages identified by unsupervised clustering. **e,** Dot plot showing canonical marker gene expression used to annotate fibroblasts, endothelial cells, smooth muscle cells, and pericytes. Dot color represents scaled average expression, and dot size represents the proportion of cells expressing each gene. **f,** Stacked bar plots showing the proportional distribution of major stromal cell types across cancer and tissue types, grouped by sample origin: tumor (T), adjacent normal (Adj.N), and healthy tissue (H). **g,** UMAP projection of reclustered fibroblasts showing eight fibroblast subpopulations. **h,** UMAP projection of fibroblasts colored by sample origin. **i,** Heatmap showing enrichment of fibroblast subtypes across tumor, adjacent normal, and healthy tissues, quantified by observed-to-expected ratio (Ro/e) analysis. **j,** Heatmap showing representative differentially expressed genes across fibroblast subtypes. **k,** Dot plot showing Reactome pathway enrichment analysis based on the top 200 differentially expressed genes from each fibroblast subtype. Dot color represents the false discovery rate (FDR), and dot size represents the number of genes associated with each pathway term. DEGs, differentially expressed genes; FDR, false discovery rate.

### Integrative single-cell analysis reveals distinct transcriptional and epigenetic programs in CAFs and NAFs

To define the molecular differences between cancer-associated fibroblasts (CAFs) and normal-activated fibroblasts (NAFs), we compared their gene-expression profiles (Fig. 2a; Supplementary Table 3). CAFs were enriched for canonical activation and matrix-remodeling genes, including POSTN, ACTA2, FAP, COL11A1, and MMP11. In contrast, NAFs showed higher expression of PLA2G2A, CLEC3B, and CFD, consistent with a more tissue-associated stromal program. This separation in marker-gene expression supports the presence of distinct tumor-associated and adjacent-normal-associated fibroblast programs.

**Figure 2.**
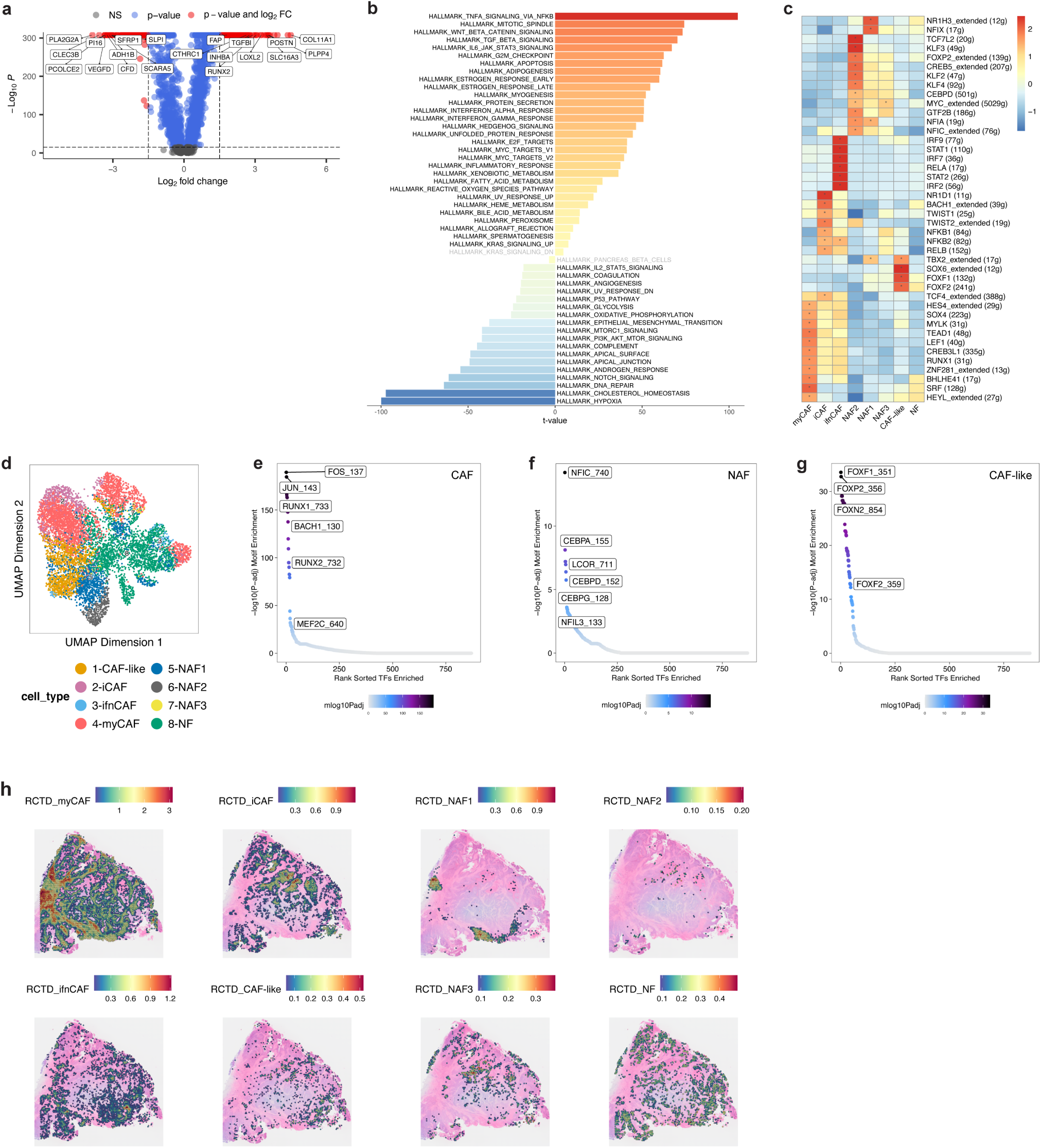
Distinct transcriptional, regulatory, and epigenetic programs define CAF and NAF heterogeneity. **a,** Volcano plot showing differentially expressed genes between CAFs and NAFs. Differential expression was performed using Seurat Find Markers with the Wilcoxon rank-sum test, considering genes expressed in at least 25% of cells in either comparison group. Positive average log_2_ fold-change values indicate genes enriched in CAFs, whereas negative values indicate genes enriched in NAFs. The y-axis shows statistical significance as −log_10_ P value. Differentially expressed genes were visualized using Enhanced Volcano with thresholds of P < 1 × 10^-15 and log_2_FC > 1.5. Selected genes are labeled. **b,** Bar plot showing differential hallmark pathway enrichment between CAFs and NAFs. Positive t-values indicate pathways preferentially enriched in CAFs, whereas negative t-values indicate pathways preferentially enriched in NAFs. **c,** Heatmap showing transcription factor regulon activity inferred by SCENIC across fibroblast subpopulations. **d,** UMAP projection of colorectal cancer snATAC-seq fibroblast cells after integration with the scRNA-seq reference, colored by annotated fibroblast subtype. **e–g,** Rank-ordered motif enrichment plots showing transcription factor motifs enriched in accessible chromatin regions associated with **e,** CAFs, **f,** NAFs, and **g,** CAF-like fibroblasts. The x-axis indicates the rank of tested transcription factors, and the y-axis represents −log_10_ adjusted P value. Top-ranked candidate transcription factors are labeled. **h,** Spatial distribution of the eight fibroblast subtype signatures in colorectal cancer tissue, estimated using Robust Cell Type Decomposition (RCTD). Color intensity represents the predicted relative abundance of each fibroblast subtype signature at each spatial location.

We next examined whether these transcriptional differences were reflected at the pathway level. Hallmark pathway enrichment analysis revealed divergent functional programs between CAFs and NAFs (Fig. 2b). Pathways with positive enrichment scores were preferentially associated with CAFs and included TNFα signaling via NF-κB, Wnt/β-catenin signaling, TGF-β signaling, and IL-6/JAK/STAT3 signaling. These pathways are consistent with inflammatory activation, stromal remodeling, and tumor-associated fibroblast functions. In contrast, pathways with negative enrichment scores were preferentially associated with NAFs and included hypoxia, cholesterol homeostasis, Notch signaling, and complement cascade-related programs. Together, these findings suggest a shift from tissue-associated fibroblast programs in NAFs toward inflammatory and tumor-associated stromal programs in CAFs.

To investigate the regulatory programs underlying fibroblast heterogeneity, we applied Single-Cell Regulatory Network Inference and Clustering (SCENIC) to infer transcription factor regulon activity across fibroblast subtypes (Fig. 2c). SCENIC analysis identified distinct regulon activity patterns among the fibroblast populations. myCAFs showed increased activity of regulons associated with RUNX1, SOX4, and TEAD1, consistent with their activated and matrix-remodeling phenotype^28,50,51^. In addition, myCAFs showed elevated expression of MYLK, a regulator of cellular contractility, adhesion, and migration, further supporting their myofibroblastic identity^52^. Because epithelial–mesenchymal transition (EMT) is linked to cancer cell plasticity and metastatic progression, we also examined the expression of the canonical EMT regulators TWIST1 and TWIST2. Both factors were increased in CAFs and are known to transcriptionally repress E-cadherin (CDH1), linking CAF-associated transcriptional programs to EMT-related pathways^53,54^. Other fibroblast subsets showed distinct regulatory features. iCAFs were enriched for NF-κB family regulons, including NFKB1, NFKB2, and RELB, consistent with their inflammatory and immunomodulatory phenotype. ifnCAFs showed strong enrichment of interferon-associated regulons, including IRF1, IRF2, IRF7, IRF9, STAT1, and STAT2, supporting their interferon-response identity. In contrast, NAFs, particularly the NAF2 subpopulation, showed increased activity of KLF family transcription factors, including KLF2, KLF3, and KLF4. These factors are associated with tissue stability and restrained proliferative states; KLF2 has been described as a tumor-suppressive regulator, KLF3 as a transcriptional repressor of pro-growth genes, and KLF4 as an important regulator of cellular homeostasis^31,32^. The coordinated enrichment of these KLF family members in NAFs supports their association with a more stable, non-tumor-associated fibroblast program. To complement the regulon activity inferred from scRNA-seq, we analyzed independent single-nucleus ATAC-seq (snATAC-seq) data from colorectal cancer tissue. The snATAC-seq data were integrated with the scRNA-seq fibroblast reference, allowing chromatin accessibility profiles to be linked with transcriptionally defined fibroblast states. After label transfer, snATAC-seq fibroblast cells were annotated according to the corresponding scRNA-seq-derived fibroblast subtypes (Fig. 2d). Motif enrichment analysis of subtype-associated accessible chromatin regions revealed distinct regulatory landscapes for CAFs, NAFs, and CAF-like fibroblasts. CAF-associated accessible peaks were enriched for RUNX1, JUN/FOS, and BACH1 motifs, whereas NAF-associated peaks were enriched for NFIC, CEBP, and LCOR motifs (Fig. 2e, f; Supplementary Table 4). Notably, RUNX1 and NFIC were identified by both SCENIC regulon analysis and snATAC-seq motif enrichment, supporting their roles as candidate regulators of CAF and NAF identity, respectively. CAF-like fibroblast-associated accessible regions showed strong enrichment for FOXF1 and FOXF2 motifs (Fig. 2g), suggesting that this population is associated with a distinct FOXF1/FOXF2-linked regulatory program. Finally, we asked whether the fibroblast programs identified by scRNA-seq could be detected within tissue space. We analyzed independent spatial transcriptomics datasets from colorectal and lung cancers using RCTD-based deconvolution (Fig. 2h; Supplementary Fig. 2a). Fibroblast subtype signatures showed non-uniform spatial patterns across tissue sections, with cancer-associated fibroblast signatures detected in localized stromal regions. Because quantitative spatial comparisons were not performed, these results were interpreted cautiously. Overall, the spatial analysis supports the tissue-level detectability of the fibroblast programs identified by single-cell analysis and provides an additional view of fibroblast heterogeneity within the tumor microenvironment. Together, these results suggest that CAFs, NAFs, and CAF-like fibroblasts represent distinct stromal states with different transcriptional and regulatory programs. In particular, CAF-like fibroblasts show a FOXF1/FOXF2-associated program, supporting their identity as a distinct fibroblast state within the tumor microenvironment.

### FOXF1/FOXF2-associated chromatin accessibility defines the CAF-like fibroblast state

To determine whether CAF-like fibroblasts are associated with a distinct transcriptional and chromatin regulatory profile, we examined FOXF1 and FOXF2 activity in this population. FOXF1 and FOXF2 showed preferential expression in CAF-like fibroblasts compared with other fibroblast subtypes (Fig. 3a, b). This expression pattern suggested that CAF-like fibroblasts may be associated with a FOXF1/FOXF2-linked regulatory program.

**Figure 3.**
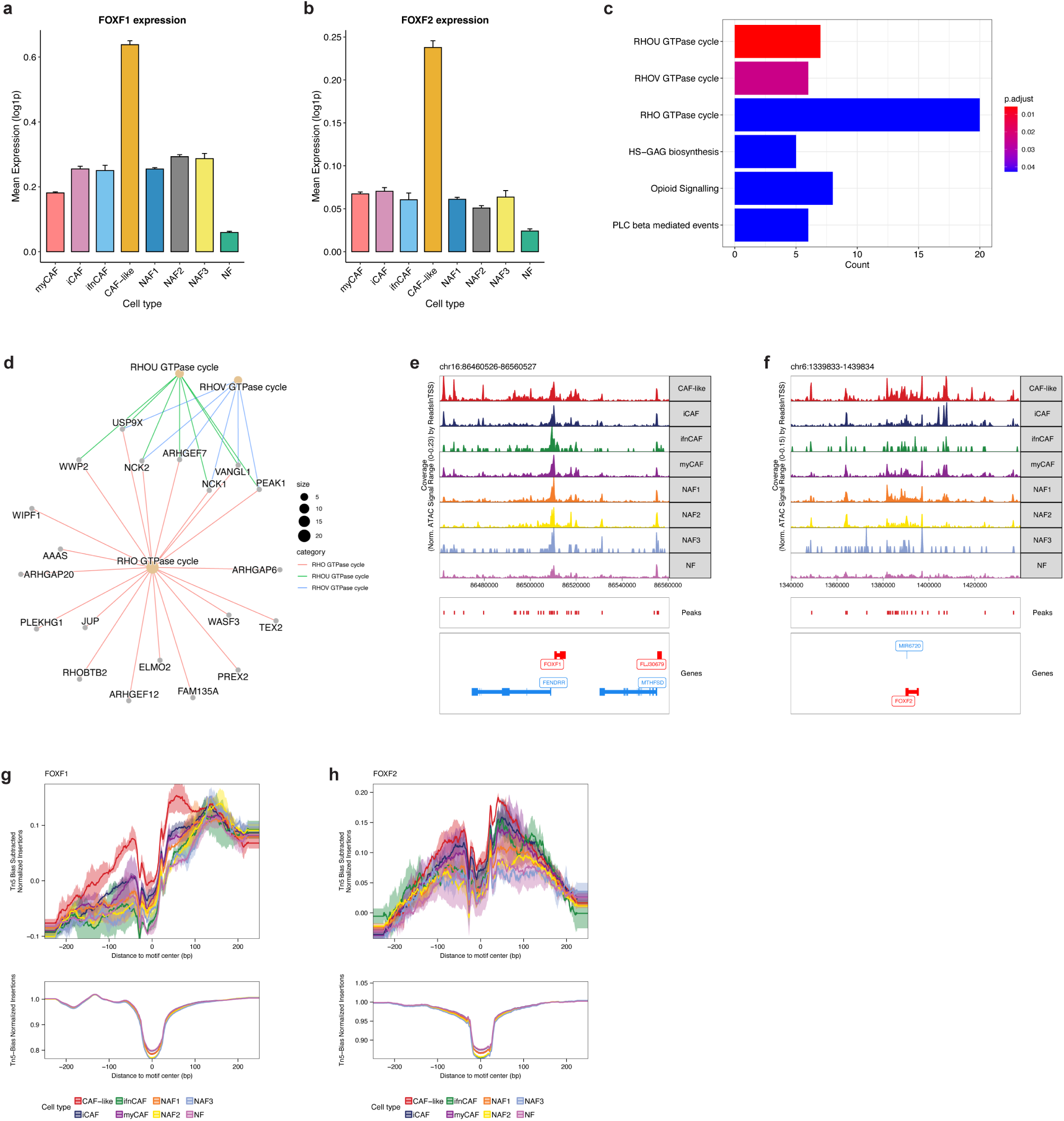
FOXF1/FOXF2-associated chromatin accessibility defines the CAF-like fibroblast state. **a, b,** Expression of FOXF1 and FOXF2 across fibroblast subtypes, showing preferential expression in CAF-like fibroblasts. **c,** Pathway enrichment analysis of combined FOXF1/FOXF2 regulon target genes. **d,** Gene-network visualization showing the relationship between FOXF1/FOXF2 regulon target genes and enriched pathways. **e, f,** Genome browser tracks showing chromatin accessibility around the FOXF1 and FOXF2 loci across fibroblast subtypes. **g,** Subtracted, normalized, and smoothed ATAC-seq footprint profile centered on FOXF1 motifs across fibroblast subtypes. The x-axis represents distance from the FOXF1 motif center, and the y-axis represents normalized chromatin accessibility signal. CAF-like fibroblasts show higher flanking accessibility and a more pronounced central depletion at the motif center, consistent with increased FOXF1 motif-associated engagement. **h,** Subtracted, normalized, and smoothed ATAC-seq footprint profile centered on FOXF2 motifs across fibroblast subtypes. The x-axis represents distance from the FOXF2 motif center, and the y-axis represents normalized chromatin accessibility signal. CAF-like fibroblasts show higher flanking accessibility and a more pronounced central depletion at the motif center, consistent with increased FOXF2 motif-associated engagement.

To further characterize the transcriptional programs associated with FOXF1/FOXF2 activity, we examined the target genes comprising the FOXF1 and FOXF2 regulons. SCENIC-inferred regulon comparison identified 53 shared target genes between FOXF1 and FOXF2, suggesting convergence on a common transcriptional program (Supplementary Fig. 2b; Supplementary Table 5). Functional enrichment analysis of the combined regulon targets revealed predominant enrichment of RHO-family GTPase signaling pathways, including RHOU- and RHOV-associated signaling, as well as broader RHO GTPase regulatory pathways (Fig. 3c, d). These pathways are linked to cytoskeletal remodeling, cell migration, and Mechan transduction during fibroblast activation, supporting the interpretation that the FOXF1/FOXF2-associated program may contribute to cellular plasticity and stromal remodeling in CAF-like fibroblasts^55,56^.

We next investigated whether the FOXF1/FOXF2-associated transcriptional program was supported at the chromatin level. Genome browser visualization around the FOXF1 and FOXF2 loci showed accessible chromatin signals in fibroblast populations, indicating regulatory accessibility around these regions (Fig. 3e, f). In addition, motif-centered footprint analysis was performed using subtracted, normalized, and smoothed ATAC-seq signal around FOXF1 and FOXF2 motifs. CAF-like fibroblasts showed the highest chromatin accessibility in the flanking regions surrounding these motifs, indicating that chromatin around FOXF1/FOXF2 motif sites is more open in the CAF-like state than in other fibroblast subtypes (Fig. 3g, h).

Importantly, CAF-like fibroblasts also displayed the most pronounced central depletion at the FOXF1 and FOXF2 motif centers, together with higher overall footprint amplitude (Fig. 3g, h). This profile is consistent with increased FOXF1/FOXF2 motif-associated engagement in CAF-like fibroblasts. In contrast, other fibroblast subtypes showed flatter and lower-amplitude footprint profiles, suggesting lower baseline accessibility and weaker evidence of FOXF1/FOXF2 motif-associated engagement. Together with the preferential expression of FOXF1 and FOXF2 and the enrichment of FOXF1/FOXF2 motifs in CAF-like-associated accessible regions, these findings support a FOXF1/FOXF2-associated chromatin regulatory program as a defining feature of the CAF-like fibroblast state.

### Pseudo time trajectory analysis identifies a transitional CAF-like state during fibroblast differentiation

To investigate how fibroblast states are related to one another during stromal activation, we performed pseudo time trajectory analysis to order fibroblasts along inferred differentiation paths. Because inactivated resident normal fibroblasts (NFs) are expected to represent the baseline stromal state, the NF-enriched region was used as the trajectory root. Using complementary trajectory approaches, including Monocle 2, Monocle 3, and Slingshot, fibroblasts were resolved into two major branches: one associated with adjacent-normal fibroblast states and the other associated with tumor-associated CAF states (Fig. 4a; Supplementary Fig. 3a, b).

**Figure 4.**
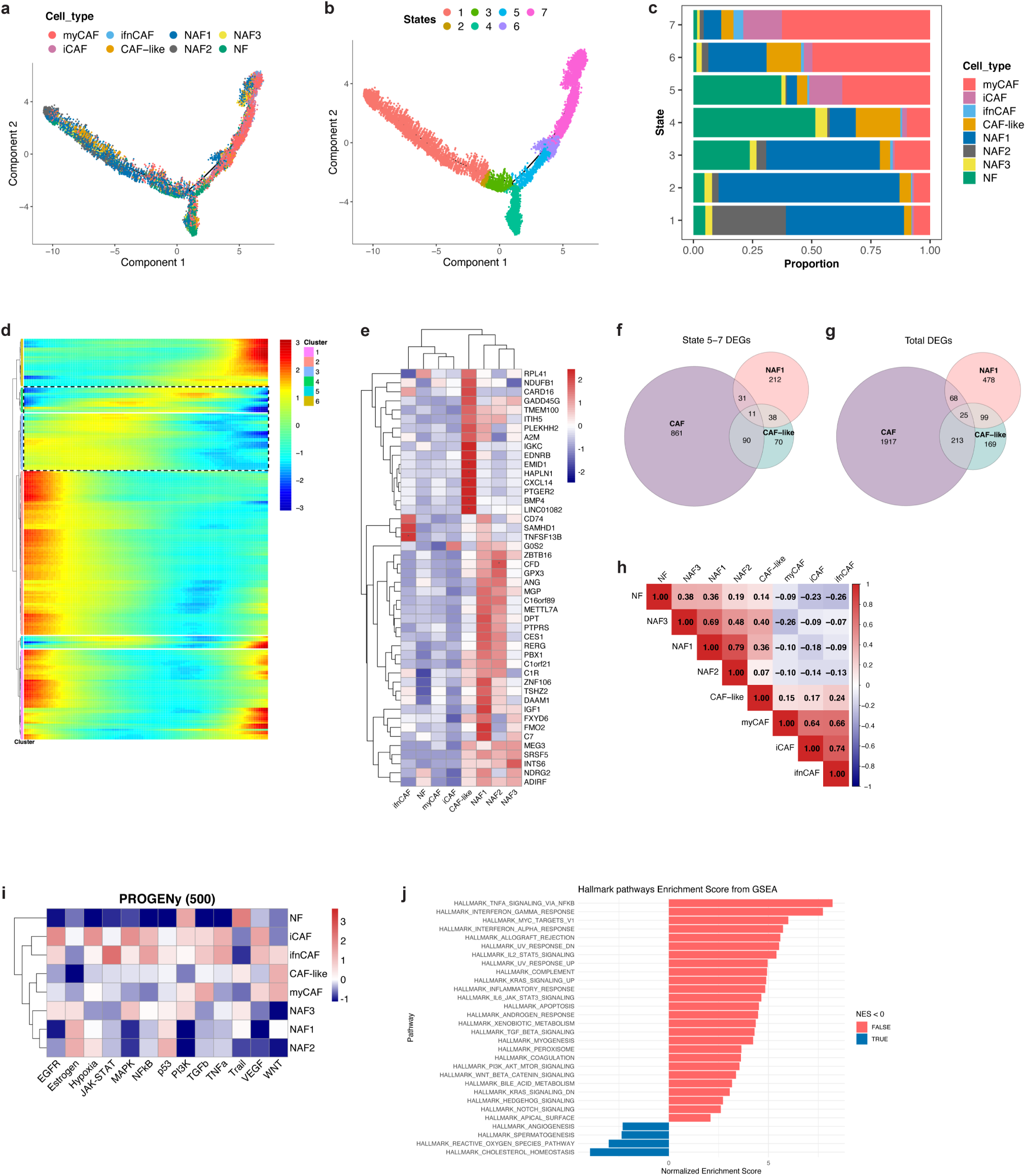
Pseudotime trajectory analysis identifies a transitional CAF-like fibroblast state. **a,** Monocle 2 pseudotime trajectory of fibroblast populations projected onto DDRTree components and colored by fibroblast subtype. The NF-enriched region was used as the trajectory root, based on the assumption that unactivated resident fibroblasts represent the baseline state. **b,** The same Monocle 2 trajectory colored by assigned cell state. State 4 corresponds to the NF-enriched root region, states 1–3 represent the adjacent-normal/NAF-associated branch, and states 5–7 represent the tumor-associated CAF branch. **c,** Relative proportion of fibroblast subtypes within each Monocle 2-defined state. **d,** Pseudotime heatmap showing expression dynamics of NAF1- and CAF-like-associated genes along the tumor-associated branch, corresponding to states 5–7. **e,** Gene expression heatmap of intermediate pseudotime gene clusters, including clusters 3 and 4, across fibroblast subtypes. **f,** Venn diagram showing the overlap among state 5–7-associated genes in NAF1 cells, CAF-like fibroblasts, and the combined CAF population comprising myCAFs, iCAFs, and ifnCAFs. **g,** Venn diagram showing the overlap among the complete differentially expressed gene profiles of NAF1 cells, CAF-like fibroblasts, and the combined CAF population comprising myCAFs, iCAFs, and ifnCAFs. **h,** Correlation matrix based on the average expression of the top 500 differentially expressed genes across fibroblast subtypes. **i,** Heatmap showing PROGENy pathway activity scores across fibroblast subtypes. **j,** Bar plot showing Normalized Enrichment Scores (NES) from Gene Set Enrichment Analysis (GSEA) of MSigDB Hallmark pathways in the CAF-like subpopulation.

In the Monocle 2 trajectory, fibroblasts were assigned to seven distinct states (Fig. 4b). State 4 was predominantly enriched for NFs and represented the inferred starting region. States 1–3 were mainly enriched for NAF populations, corresponding to the adjacent non-cancerous fibroblast branch. In contrast, states 5–7 contained a higher proportion of CAF populations, indicating the tumor-associated branch (Fig. 4c). Notably, NAF1 and CAF-like fibroblasts were also enriched within states 5 and 6 of this tumor-associated branch. Together with the Slingshot trajectory, which positioned NAF1 and CAF-like fibroblasts along the CAF-directed lineage, these results suggest that CAF-like fibroblasts may occupy an intermediate position during the transition from NF-like states toward CAF-associated states (Supplementary Fig. 3b).

To further examine this putative transitional phase, we focused on cells within the tumor-associated branch, corresponding to Monocle 2 states 5–7. By intersecting genes expressed within this branch with fibroblast subtype-specific differentially expressed genes, we identified NAF1-associated and CAF-like-associated gene sets within states 5–7 (Supplementary Tables 6 and 7). We then visualized the expression dynamics of these genes along pseudo time using heatmap analysis (Fig. 4d; Supplementary Fig. 4). This analysis identified intermediate gene clusters, particularly clusters 3 and 4, that showed elevated expression during the mid-phase of pseudo time and were more strongly enriched in CAF-like fibroblasts than in NAF1 cells (Fig. 4e). These expression dynamics support the placement of CAF-like fibroblasts within an intermediate transcriptional phase of the tumor-associated trajectory.

We next asked whether CAF-like fibroblasts share gene programs with CAFs within the tumor-associated branch. Comparison of state 5–7-associated genes among NAF1 cells, CAF-like fibroblasts, and the combined CAF population comprising myCAFs, iCAFs, and ifnCAFs showed substantial overlap between CAF-like and CAF gene signatures. Among the 209 CAF-like state 5–7-enriched genes, 101 were shared with the combined CAF population (Fig. 4f; Supplementary Table 8). This overlap suggests that CAF-like fibroblasts carry a considerable part of the CAF-associated transcriptional program within the tumor-associated trajectory branch, while still retaining a distinct gene-expression profile.

To determine whether this relationship was also evident beyond the pseudo time-defined branch, we compared the complete differentially expressed gene profiles of CAF-like fibroblasts, NAF1 cells, and the combined CAF population. Consistent with the branch-specific analysis, CAF-like fibroblasts showed a greater overlap with CAFs than with NAF1 cells, with 238 of 506 CAF-like DEGs shared with the combined CAF population (Fig. 4g). Correlation analysis based on the average expression of the top 500 DEGs further showed that CAF-like fibroblasts were more positively correlated with CAF populations than with NAF subtypes (Fig. 4h). These results indicate that CAF-like fibroblasts are transcriptionally closer to CAFs than to NAFs, while still maintaining a distinct identity.

Finally, we examined pathway activity across fibroblast subtypes. PROGENy-based pathway activity analysis identified WNT signaling as a prominently active pathway in both CAF-like fibroblasts and myCAFs (Fig. 4i). This observation was further supported by Gene Set Enrichment Analysis, which showed enrichment of WNT signaling within the CAF-like cluster (Fig. 4j; Supplementary Fig. 3c). Collectively, these findings support CAF-like fibroblasts as a distinct, putative transitional state within the inferred trajectory from NF-like fibroblasts toward CAF-associated states and suggest that WNT signaling may be linked to this transition.

### Non-canonical WNT and PAR-associated communication programs define the CAF-like fibroblast state

Building on pathway analyses showing WNT pathway activity in CAF-like fibroblasts (Fig. 4i, j), we next examined the WNT signaling components associated with this transitional state. WNT ligands were grouped into canonical and non-canonical WNT classes based on established classification^46^. Within CAF-like fibroblasts, non-canonical WNT activity was significantly higher than canonical WNT activity, as reflected by increased ncWNT module scores (Fig. 5a). Across fibroblast subtypes, ncWNT activity was increased in CAF-like fibroblasts compared with NAF and NF populations, supporting preferential association of non-canonical WNT signaling with the CAF-like/tumor-associated fibroblast program (Fig. 5b). Among individual WNT ligands, WNT5A was expressed in CAF-like fibroblasts and other activated fibroblast states, while showing lower expression in NAF and NF populations (Fig. 5c). Given the reported roles of WNT5A in chronic inflammation and stromal remodeling^57^, these findings suggest that WNT5A-associated non-canonical WNT signaling is a prominent feature of CAF-like fibroblast activation. TCGA bulk RNA-seq analysis further showed variable WNT5A expression between tumor and adjacent normal tissues across cancer types, indicating context-dependent regulation of WNT5A expression (Supplementary Fig. 5a). To determine whether CAF-like fibroblasts participate in intercellular communication within the tumor microenvironment, we performed Cell Chat analysis using LC and CRC datasets. Global signaling analysis showed that CAF-like fibroblasts displayed both outgoing and incoming communication programs, suggesting that this population can actively send and receive signals within the stromal microenvironment (Fig. 5d; Supplementary Fig. 5b). Several pathways were predicted to involve CAF-like fibroblasts, including HGF, PDGF, PAR, and ncWNT signaling. Because multiple pathways were detected, we next examined ligand–receptor contribution patterns to identify the most interpretable communication axes for downstream analysis. This analysis highlighted GZMA–F2R as the dominant ligand–receptor pair, followed by WNT5A–MCAM, providing a rationale for focusing on PAR- and ncWNT-associated signaling programs (Fig. 5e; Supplementary Fig. 5c). PAR signaling was mainly represented by T cell–derived GZMA interacting with F2R on CAF-like fibroblasts. Network centrality analysis indicated that T cells acted as major senders, whereas CAF-like fibroblasts showed receiver activity within the PAR signaling network (Fig. 5f; Supplementary Fig. 5d). This suggests a candidate immune–stromal communication axis in which cytotoxic immune-cell-derived GZMA may signal to F2R-expressing CAF-like fibroblasts. In parallel, ncWNT signaling showed sender activity in CAF-like and related activated fibroblast populations, with WNT5A–MCAM contributing to communication with endothelial or stromal compartments (Fig. 5g). Thus, PAR signaling highlighted a potential immune–fibroblast interaction, whereas ncWNT signaling highlighted a WNT5A-associated stromal communication program. To assess whether the predicted GZMA–F2R interaction was spatially plausible, we analyzed independent LUAD and CRC spatial transcriptomics datasets. Spatial mapping showed tissue-level proximity between GZMA and F2R expression, supporting possible co-occurrence of this ligand–receptor pair within the tumor architecture (Fig. 5h; Supplementary Fig. 5e). Although spatial proximity does not prove direct signaling, it supports the biological plausibility of the CellChat-predicted GZMA–F2R interaction. GZMA is secreted by cytotoxic immune cells and has been reported to promote tumor suppression and T cell–mediated cytotoxicity through JAK2/STAT1 signaling^58^; however, the functional consequences of GZMA–F2R signaling in fibroblasts remain largely unexplored. We next evaluated the clinical relevance of the CAF-like fibroblast signature across TCGA cohorts. Kaplan–Meier analysis of TCGA-STAD showed that patients with high CAF-like signature expression had poorer overall survival than patients with low CAF-like signature expression (Fig. 5i). However, survival analysis across additional TCGA cohorts showed that the prognostic association of the CAF-like signature was not uniform across cancer types (Supplementary Fig. 6a–f). For example, high CAF-like signature expression was associated with poorer survival in TCGA-ESCA, whereas other cohorts showed neutral or non-significant associations. These findings indicate that the CAF-like signature should not be interpreted as a universal poor-prognosis marker, but rather as a context-dependent stromal program. To further examine the STAD-specific association, TCGA-STAD samples were stratified into tumor and adjacent normal tissue compartments. In tumor samples, high CAF-like signature expression was associated with reduced overall survival, whereas adjacent normal samples did not show a statistically significant adverse association (Supplementary Fig. 6g, h). Univariate Cox proportional hazards analysis further identified F2R and EDNRB as CAF-like signature genes associated with increased mortality risk in the overall STAD cohort (Fig. 5j). Tissue-stratified Cox analysis showed that these risk associations were mainly observed in tumor tissue, while adjacent normal tissue showed no significant association for individual CAF-like genes (Supplementary Fig. 6i). Notably, F2R was also the receptor component of the predicted GZMA–F2R interaction, linking the cell–cell communication analysis with the tumor-specific clinical relevance of the CAF-like state. Together, these findings suggest that CAF-like fibroblasts are characterized by non-canonical WNT activity, WNT5A-associated stromal signaling, and a candidate immune–stromal GZMA–F2R/PAR communication axis. Clinically, the CAF-like signature shows context-dependent prognostic relevance, with stronger adverse association in the tumor compartment of TCGA-STAD.

**Figure 5.**
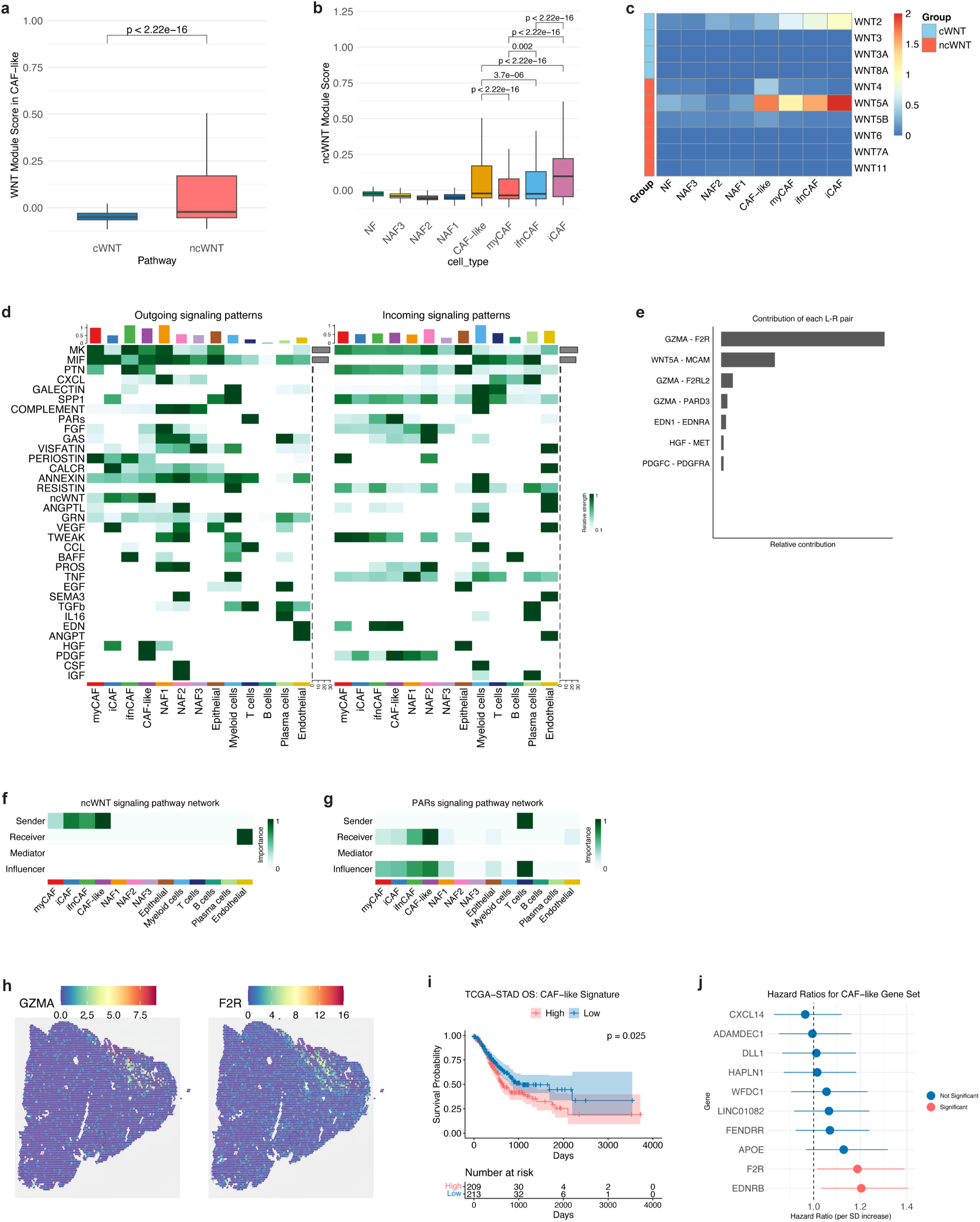
Non-canonical WNT activity, PAR-associated communication, and clinical relevance of CAF-like fibroblasts. **a,** Box plot comparing canonical WNT (cWNT) and non-canonical WNT (ncWNT) module scores within CAF-like fibroblasts. **b,** Box plot showing ncWNT module scores across fibroblast subtypes. **c,** Heatmap showing expression of canonical and non-canonical WNT ligands across fibroblast subtypes. **d,** CellChat analysis of lung cancer showing outgoing and incoming signaling patterns across fibroblast subclusters and major tumor microenvironment cell types. **e,** Ligand–receptor contribution analysis showing prominent interaction pairs associated with CAF-like communication programs, including GZMA–F2R and WNT5A–MCAM. **f,** Network centrality heatmap for PAR signaling in lung cancer, showing sender, receiver, mediator, and influencer activities across cell types. **g,** Network centrality heatmap for ncWNT signaling in lung cancer, showing sender, receiver, mediator, and influencer activities across cell types. **h,** Spatial transcriptomics plots from LUAD tissues showing spatial expression patterns of GZMA and its receptor F2R. **i,** Kaplan–Meier overall survival curve for TCGA-STAD patients stratified into high and low CAF-like signature groups based on the median signature score. **j,** Forest plot showing univariate Cox proportional hazards regression analysis for individual CAF-like signature genes in TCGA-STAD. Hazard ratios are shown per standard deviation increase in gene expression. Red markers indicate statistically significant associations, whereas blue markers indicate non-significant associations.

**Figure 6.**
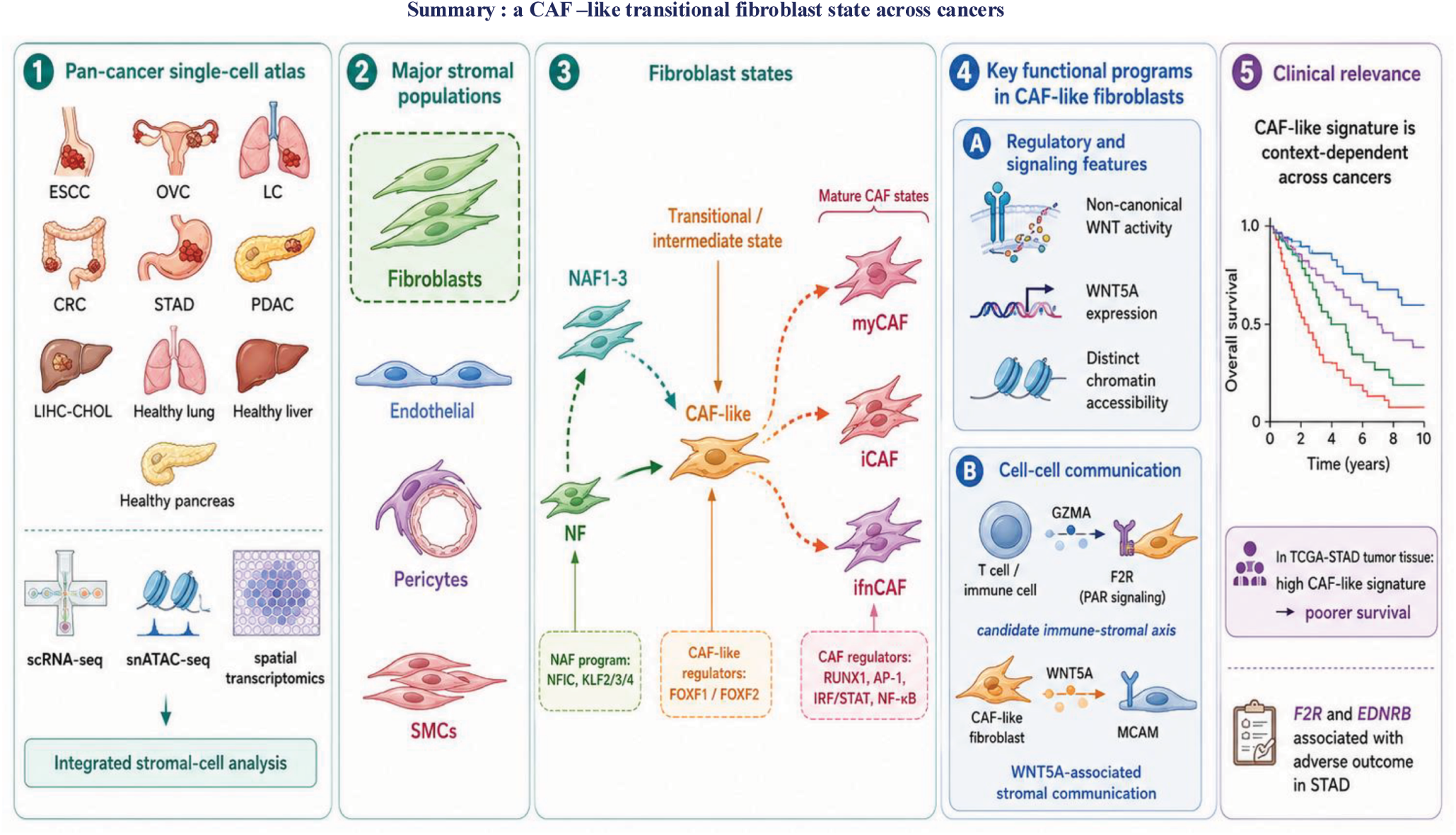
Graphical summary: a CAF-like transitional fibroblast state across cancers. CAF-like fibroblasts represent a FOXF1/FOXF2-associated transitional stromal state linking normal fibroblasts to activated CAFs, with non-canonical WNT activity, candidate GZMA–F2R immune–stromal signaling, and context-dependent clinical relevance.

## Discussion

This study identifies a CAF-like fibroblast population positioned between normal-associated fibroblast states and established CAF subsets across cancers. Rather than supporting a binary view in which fibroblasts are either normal or fully tumor-associated, our data point to a spectrum of stromal activation. Within this spectrum, the CAF-like population appears to represent a transitional state carrying features of both tissue-associated fibroblasts and activated CAFs. This reframes fibroblast heterogeneity as a dynamic process rather than a fixed set of endpoint phenotypes, highlighting intermediate states that may actively shape how the tumor stroma evolves. A key implication of these findings is that CAF-like fibroblasts may act as a molecular bridge between NAFs and mature CAF states. Because this population is detected in both adjacent normal and tumor compartments, stromal activation may begin before fibroblasts acquire fully established CAF features. This interpretation is consistent with the broader view that CAF identity is shaped by cellular origin, tissue context, cancer type, and local microenvironmental signals^22,23^. Our data extend this concept by nominating a distinct intermediate state, rather than treating CAF development simply as the gradual loss of normal fibroblast identity. Under this model, the CAF-like state may function as a poised stromal state that can be directed toward specialized CAF programs depending on the dominant signals within the tumor microenvironment. The regulatory profile of CAF-like fibroblasts supports this interpretation. FOXF1 and FOXF2 were preferentially associated with the CAF-like state at both the transcriptomic and chromatin accessibility levels, contrasting with NAFs and mature CAFs, where NFIC/KLF family members and RUNX1/AP-1-related programs marked more stable normal-associated or activated CAF identities. FOXF1 and FOXF2 have known roles in mesenchymal lineage specification and stromal development^59–61^, and their enrichment in CAF-like fibroblasts suggests that this intermediate state may be maintained by a distinct mesenchymal regulatory program. However, these data do not establish FOXF1/FOXF2 as direct drivers of CAF differentiation. Instead, they nominate FOXF1/FOXF2 as candidate regulators of fibroblast plasticity and provide a concrete, testable entry point for future functional studies. Functionally, CAF-like fibroblasts were marked by non-canonical WNT activity, particularly WNT5A-associated signaling. WNT signaling was first identified at the pathway level in the CAF-like state and was then resolved into a more specific non-canonical WNT program. This is notable given the reported roles of WNT5A in inflammation, cell migration, tissue remodeling, and context-dependent cancer biology^57^. In our data, WNT5A-associated activity was higher in CAF-like fibroblasts than in NAF and NF populations, supporting the idea that this pathway is acquired during stromal activation. However, WNT5A expression also varied across cancer types, arguing against a universal interpretation. Functional enrichment of the FOXF1/FOXF2 regulon targets further revealed predominant activity of RHO-family GTPase signaling, including RHOU and RHOV (Fig. 3c, d). This is mechanistically relevant given that non-canonical WNT5A signaling activates RHO GTPases to drive cytoskeletal remodeling in fibroblasts^62,63^, and RHOU itself is a known transcriptional target of non-canonical WNT activation^64^. Together, these findings suggest that the FOXF1/FOXF2-associated RHOU/RHOV program and CAF-like WNT5A activity may reflect a coherent signaling axis linking transcriptional regulation to cytoskeletal remodeling, rather than two independent observations, warranting future functional validation. Thus, non-canonical WNT activity is best understood as a context-sensitive feature of CAF-like activation rather than a fixed marker of tumor-promoting behavior. Cell–cell communication analysis added another layer to this model. Among the predicted signaling pathways involving CAF-like fibroblasts, ligand–receptor contribution analysis identified WNT5A–MCAM and GZMA–F2R as prominent interaction pairs. These findings point to two candidate communication programs: WNT5A-associated stromal communication and GZMA–F2R/PAR-associated immune–stromal communication. The GZMA–F2R axis is particularly interesting because it links cytotoxic immune-cell-derived signals with CAF-like fibroblasts, raising the possibility that immune activity may influence stromal states in addition to acting on tumor cells. Spatial co-occurrence of GZMA and F2R supports the tissue-level plausibility of this interaction, although it does not prove direct signaling. Therefore, the GZMA–F2R/PAR axis should be viewed as a candidate mechanism of immune–stromal crosstalk rather than an established functional pathway. The prognostic analysis adds an important caution to the interpretation of CAF-like biology. The CAF-like signature was not uniformly associated with poor outcome across cancers; instead, its survival association depended on cancer type and tissue context. This pattern is consistent with the broader understanding that CAFs and CAF-like states can have divergent effects depending on tumor type, anatomical site, immune context, and stromal composition^17,20,21^. Therefore, CAF-like fibroblasts should not be labeled as universally tumor-promoting or universally protective. Their significance lies in their plasticity and context dependence. Within this framework, the STAD findings are particularly informative. In TCGA-STAD, high CAF-like signature expression was associated with poorer survival, and this association was mainly observed in tumor tissue rather than adjacent normal tissue. This suggests that the same stromal program may carry different biological meaning depending on where it is expressed. In adjacent normal tissue, a CAF-like signature may reflect wound repair, tissue remodeling, or a poised stromal state. In tumor tissue, the same program may be coupled to malignant–stromal interactions and adverse outcome. The tumor-specific association of F2R and EDNRB with increased mortality risk further supports this interpretation. Because F2R was also part of the predicted GZMA–F2R communication axis, it provides a link between the signaling analysis and the clinical findings. Nevertheless, F2R should be considered a candidate biomarker and mechanistic lead, not yet a validated therapeutic target. Taken together, these findings support a model in which CAF-like fibroblasts represent a FOXF1/FOXF2-associated transitional stromal state with context-dependent functional and clinical consequences. This state is marked by regulatory plasticity, non-canonical WNT activity, WNT5A-associated stromal communication, and a candidate GZMA–F2R/PAR immune–stromal axis. The broader implication is that stromal-targeting strategies should avoid treating CAFs as a uniform population. Instead, therapeutic interpretation should account for fibroblast state, tissue compartment, and cancer type. Several limitations should be acknowledged. First, the inferred NAF-to-CAF-like-to-CAF transition is based on cross-sectional single-cell data and pseudo time modeling; lineage tracing will be required to confirm true cellular progression. Second, CellChat and spatial transcriptomics support predicted and spatially plausible interactions, but they do not demonstrate direct molecular signaling. Third, survival analyses based on bulk TCGA data may be influenced by tumor purity, stromal abundance, immune infiltration, and tissue composition. Future studies should test whether perturbing FOXF1/FOXF2 alters CAF-like state formation, whether WNT5A–MCAM signaling contributes to stromal remodeling, and whether GZMA–F2R/PAR signaling functionally modifies fibroblast activation in tumor tissue.

In conclusion, this study defines a CAF-like transitional fibroblast state across cancers and links it to FOXF1/FOXF2-associated regulation, non-canonical WNT signaling, and a candidate immune–stromal communication axis through GZMA–F2R/PAR. Its clinical relevance is not universal but context dependent, with a stronger adverse association in the tumor compartment of STAD. Together, these findings provide a framework for understanding CAF evolution as a dynamic, tissue-specific process and nominate concrete regulatory and communication pathways for future stromal-focused investigation.

## Data availability

The datasets analyzed in this study are publicly available. Single-cell RNA sequencing data for colorectal cancer are available from the Gene Expression Omnibus (GEO; https://www.ncbi.nlm.nih.gov/geo/) under accession numbers GSE132465, GSE132257, and GSE144735. Lung adenocarcinoma and non-small cell lung cancer datasets are available under GEO accession numbers GSE131907 and GSE148071, respectively. Liver cancer datasets, comprising hepatocellular carcinoma and intrahepatic cholangiocarcinoma, are available under GEO accession numbers GSE125449, GSE138709, and GSE142784. Pancreatic ductal adenocarcinoma datasets are available under GEO accession numbers GSE154778 and GSE155698. Esophageal squamous cell carcinoma data are available under GEO accession number GSE160269. Healthy donor datasets for lung, liver, and pancreatic tissue are available under GEO accession numbers GSE122960, GSE115469, and GSE131886, respectively. Ovarian cancer data are available from the Lambrechts Lab data portal (https://lambrechtslab.sites.vib.be/en/dataaccess). Stomach adenocarcinoma data are available from the Stanford DNA Discovery portal (https://dna-discovery.stanford.edu/research/datasets/) and from the European Nucleotide Archive (ENA) under accession number PRJEB45598. Spatial transcriptomics datasets for colorectal and lung cancer are available from the 10x Genomics website (https://www.10xgenomics.com/datasets). The snATAC-seq dataset for colorectal cancer is available from GEO under accession number GSE306459.

## Acknowledgements

This research was supported by the Bio and Medical Technology Development Program of the National Research Foundation of Korea (NRF), funded by the Ministry of Science and ICT (MSIT), Republic of Korea (Grant No. NRF-2017M3A9A7050803); the Korea Health Technology R&D Project through the Korea Health Industry Development Institute (KHIDI), funded by the Ministry of Health and Welfare, Republic of Korea (Grant No. RS-2026-25510743); and the BK21 FOUR Program of the National Research Foundation of Korea (NRF), funded by the Ministry of Education, Republic of Korea.

## Author Contributions

B.M. and A.V. wrote the manuscript. B.M. performed all bioinformatics and computational analyses and interpreted the results. T.N. assisted with the bioinformatics analyses throughout the study. Y.H.B. helped in single cell nucleus data analysis and critically reviewed the manuscript. A.V. also reviewed and edited the manuscript and provided critical comments. W.Y.P. conceived and supervised the study and provided overall scientific guidance. All authors read and approved the final manuscript.

