## Supplementary Figures and tables for "Pan-cancer single-cell analysis identifies a FOXF1/FOXF2-associated transitional CAF-like fibroblast state": Supplementary_figures_300dpi_compressed.pdf

### Supplementary Figure 1

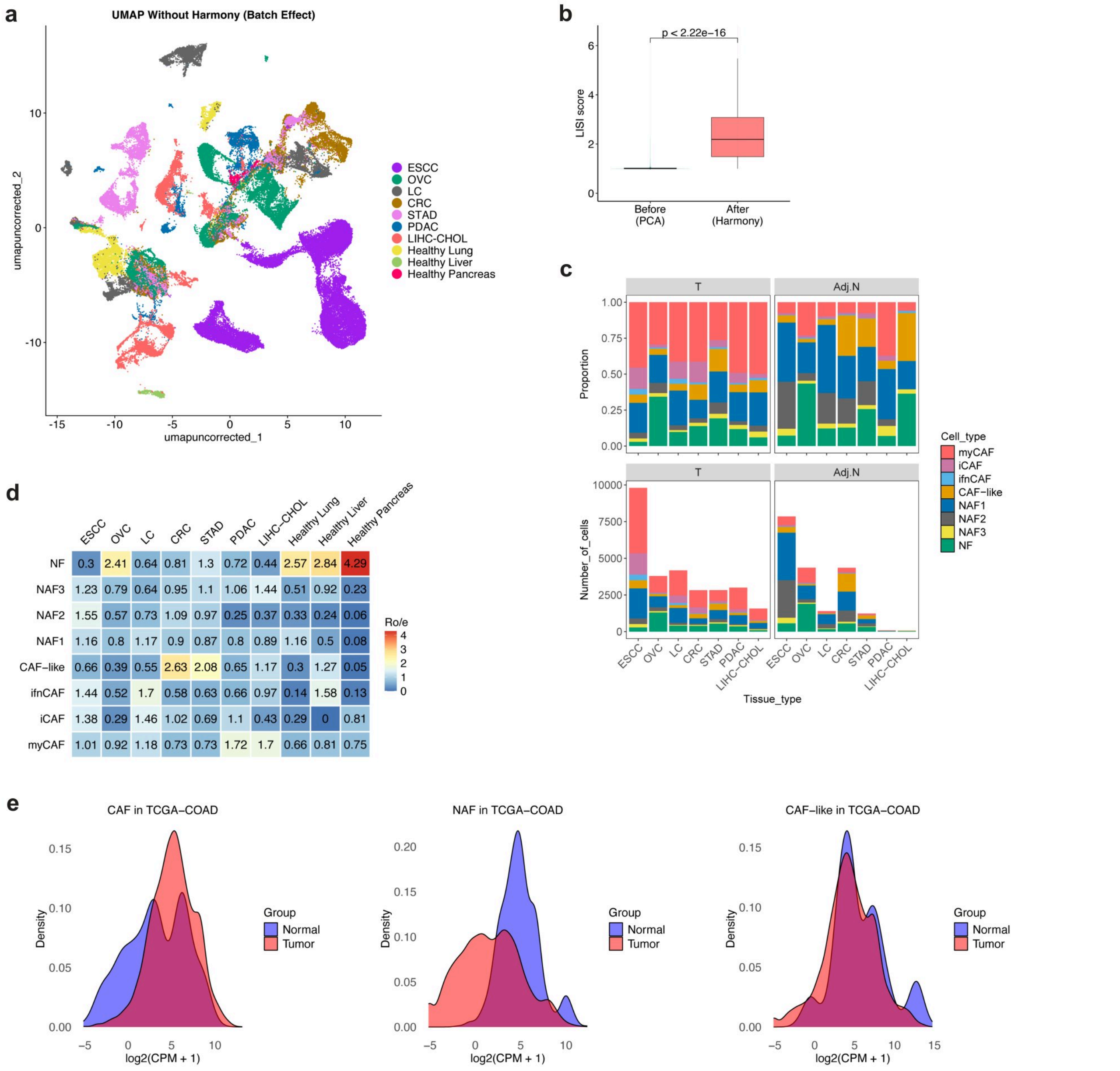


**Supplementary Figure 1. Data integration of stromal cells, distribution, and validation of fibroblast subtypes across pan-cancer cohorts. a,** UMAP projection of stromal cells before Harmony integration, colored by specific tumor types and healthy donor tissue origins. **b,** Box plot quantifying dataset integration quality using the Local Inverse Simpson's Index scored by tumor type (LISI) before and after Harmony batch correction. **c,** Bar plots illustrating the absolute cellular counts (bottom) and proportional distributions (top) of the eight identified fibroblast subpopulations across multiple cancer types. Data are stratified by tissue origin: Tumor (T, left) and Adjacent Normal (Adj.N, right). **d,** Heatmap displaying the relative enrichment of each fibroblast subtype across distinct cancer types and healthy donor tissues, quantified by observed-to-expected ratio (Ro/e) analysis. **e,** Validation of fibroblast expression signatures in bulk transcriptomic data. Density profiles depict the expression levels $log_2$(CPM + 1) of the CAF, NAF, and transitional CAF-like gene signatures within normal adjacent (blue) and tumor (red) tissues from the TCGA colon adenocarcinoma (TCGA-COAD) cohort.

### Supplementary Figure 2

**a**

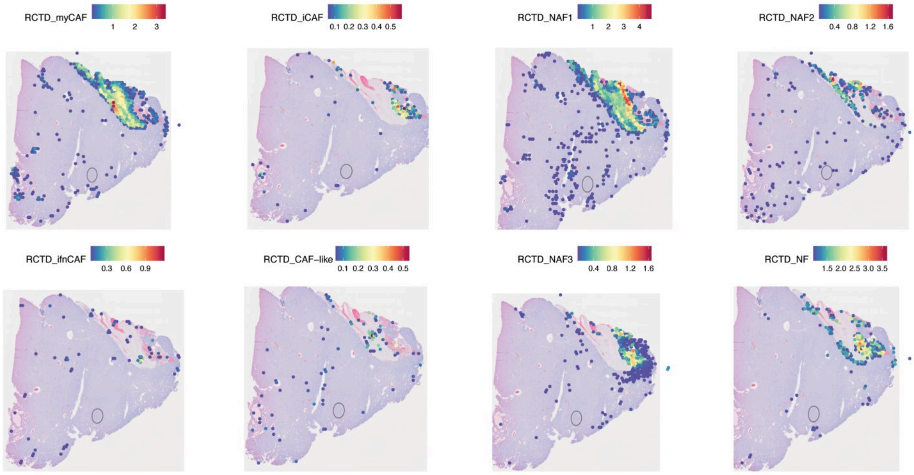


**b**

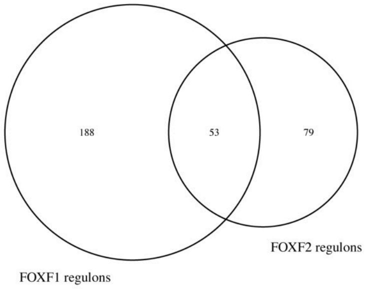


**Supplementary Figure 2.**
**a, Spatial distribution of the eight fibroblast subtype signatures in lung cancer tissue,** estimated using Robust Cell Type Decomposition (RCTD). Color intensity represents the predicted relative abundance of each fibroblast subtype signature at each spatial location.
**b, Venn diagram showing the overlap between FOXF1 and FOXF2 regulons inferred by SCENIC.** 53 genes were shared between both regulons, suggesting the presence of a common FOXF1/FOXF2 regulatory network.

#### Supplementary Figure 3

a

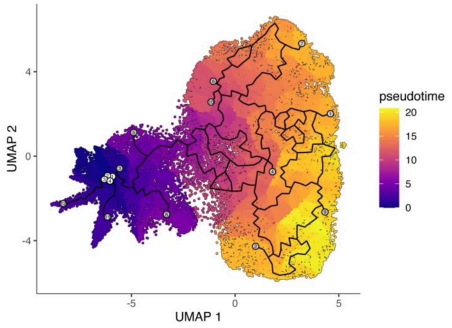


b

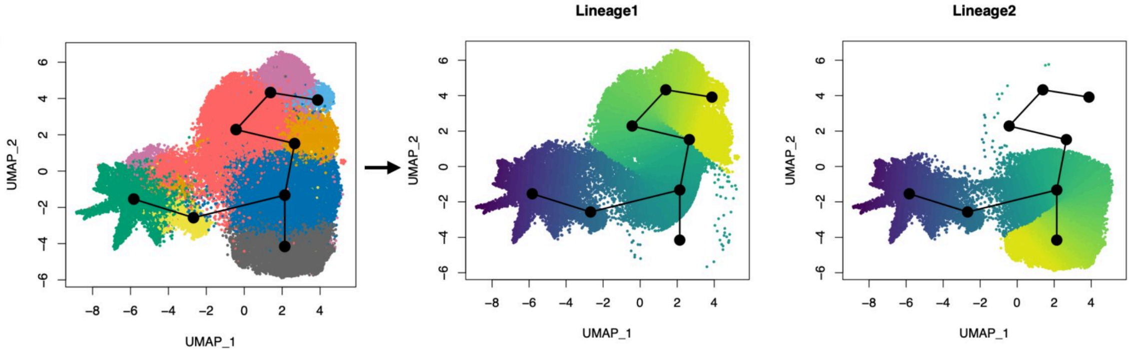


c

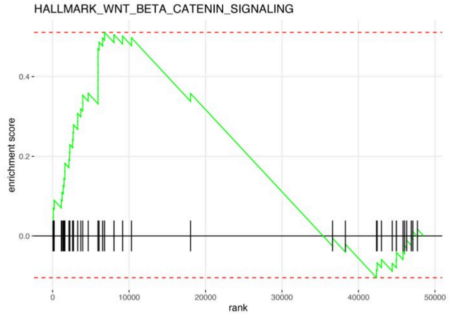


**Supplementary Figure 3. Pseudotemporal trajectory analysis of fibroblast differentiation lineages.**
**a,** Single-cell trajectory inference generated by Monocle3. The UMAP plot illustrates the continuous pan-cancer fibroblast differentiation trajectory, with cells colored according to their calculated pseudotime score along the developmental path (from dark blue [early/quiescent state] to bright yellow [late/activated state]). Black lines indicate the reconstructed principal graph structure, detailing the bifurcation points within the fibroblast population.
**b,** Alternative lineage topology reconstruction using Slingshot. Left: UMAP projection of fibroblast subpopulations, overlaid with the principal curves (black lines and nodes) rooted in the normal fibroblast (NF) cluster. Center and right: Individual deconstruction of the trajectory into two distinct developmental pathways: Lineage 1 (center) and Lineage 2 (right) with cells gradient-colored by lineage-specific pseudotime values.
**c, Gene Set Enrichment Analysis (GSEA)** showing enrichment of the HALLMARK_WNT_BETA_CATENIN_SIGNALING gene set in CAF-like fibroblasts. The green curve represents the running enrichment score, and black tick marks indicate the positions of WNT pathway genes within the ranked gene list.

### Supplementary Figure 4

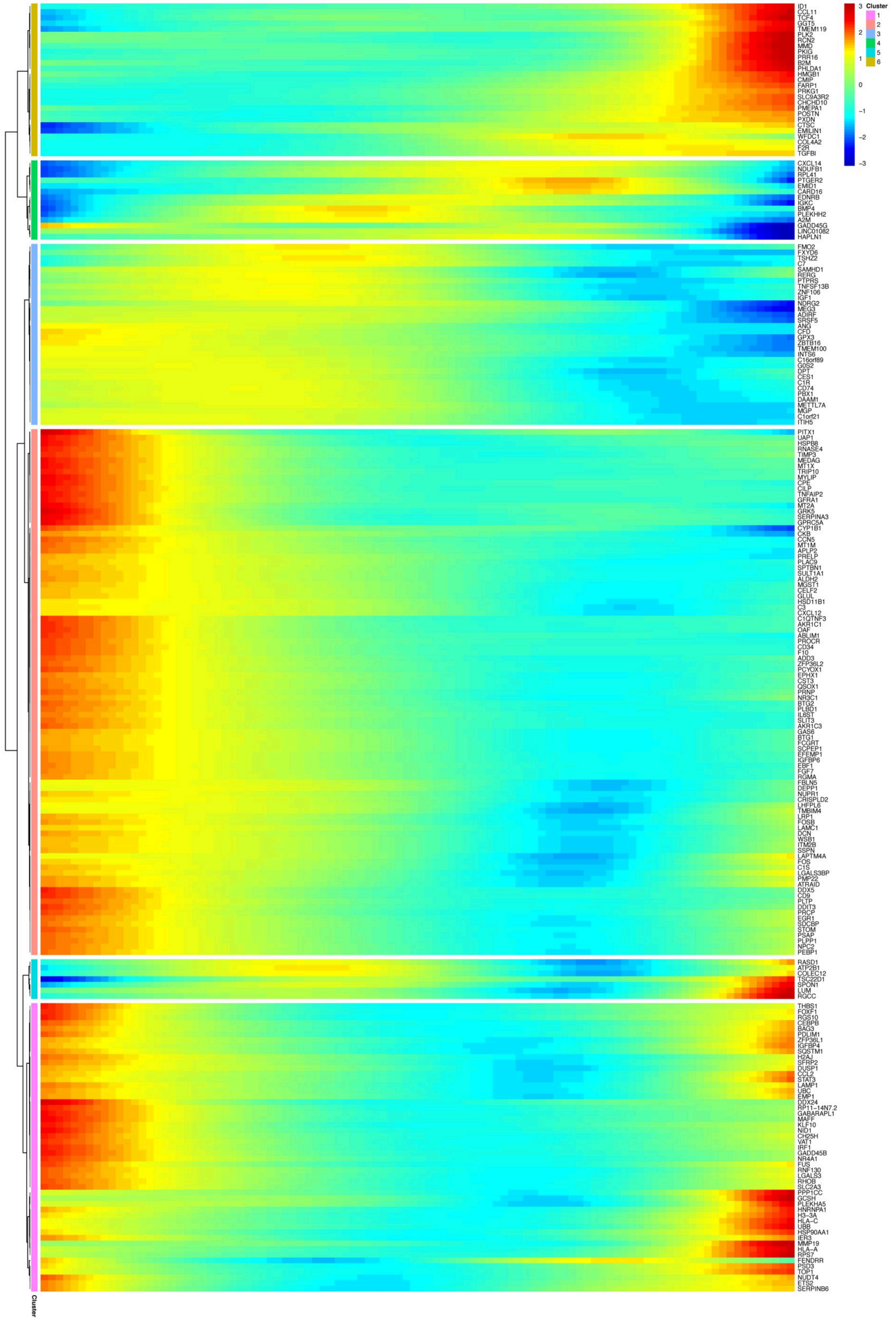


**Supplementary Figure 4. Pseudotime expression dynamics of NAF1- and CAF-like-associated genes.** Pseudotime heatmap showing expression dynamics of NAF1- and CAF-like-associated genes along the tumor-associated branch, corresponding to states 5–7.

### Supplementary Figure 5

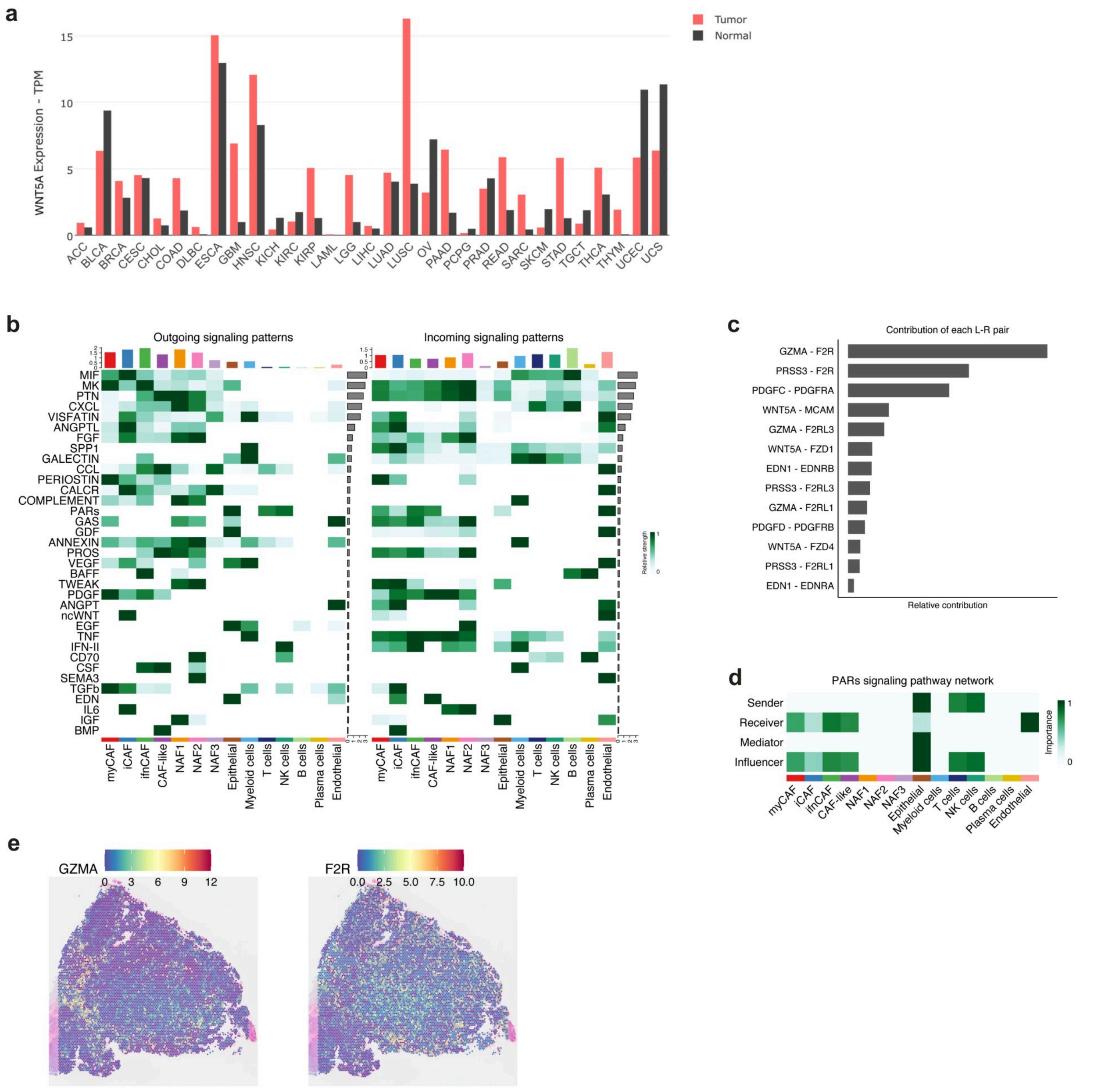


**Supplementary Figure 5. Pan-cancer WNT5A expression in TCGA and CAF-like communication programs in colorectal cancer.**

**a,** Bar chart showing the differential expression of WNT5A in tumor (red) and adjacent normal (black) tissues across multiple cancer types, generated using the GEPIA web server based on TCGA data.

**b,** CellChat analysis of colorectal cancer showing outgoing and incoming signaling patterns across fibroblast subclusters and major tumor microenvironment cell types.

**c,** Ligand–receptor contribution analysis showing prominent interaction pairs associated with CAF-like communication programs, including GZMA–F2R.

**d,** Network centrality heatmap for PAR signaling in colorectal cancer, showing sender, receiver, mediator, and influencer activities across cell types.

**e,** Spatial transcriptomics plots from colorectal cancer tissues showing spatial expression patterns of GZMA and its receptor F2R.

### Supplementary Figure 6

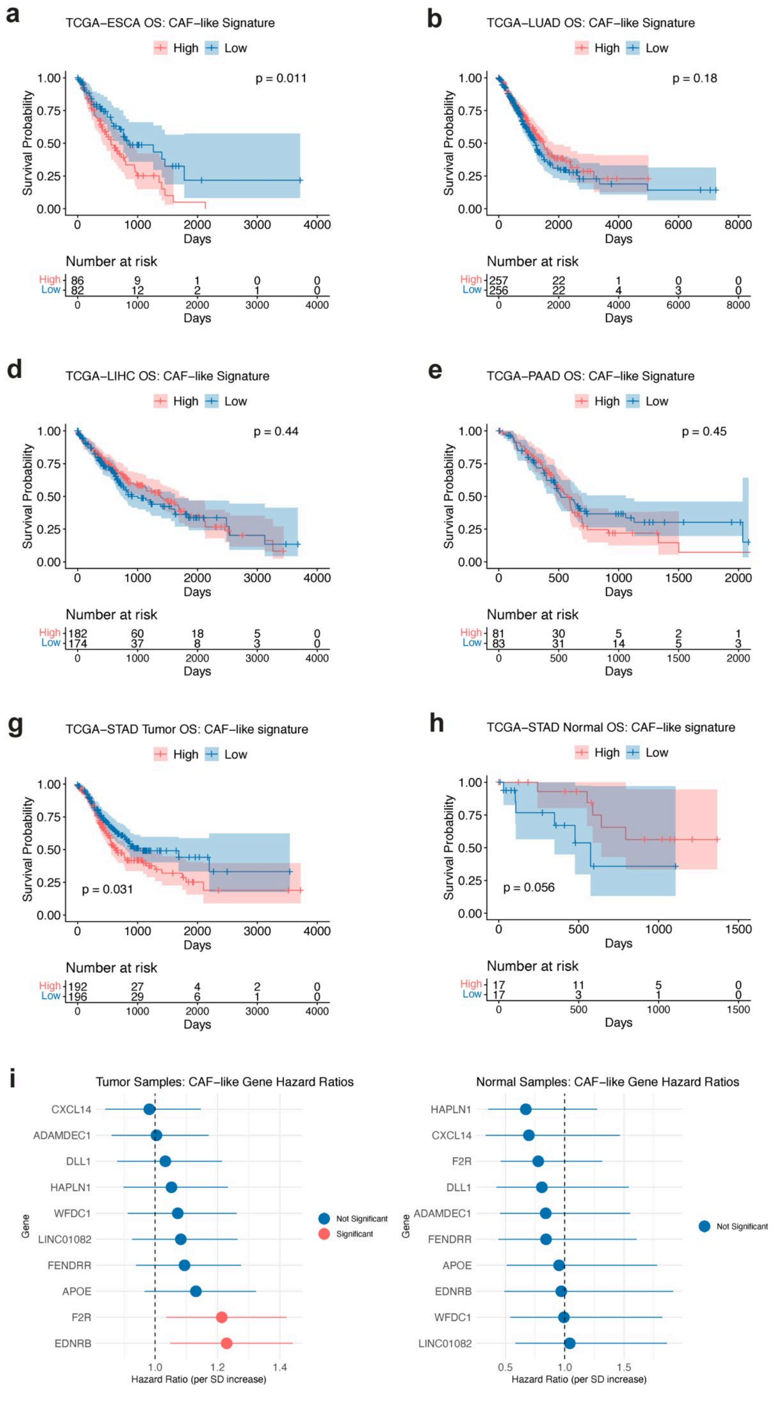


**Supplementary Figure 6. Prognostic significance of the CAF-like gene signature in TCGA cohorts.**
**a - f,** Kaplan–Meier overall survival (OS) curves for patients across TCGA-ESCA **(a)**, TCGA-LUAD **(b)**, TCGA-COAD **(c)**, TCGA-LIHC **(d)**, TCGA-PAAD **(e)**, and TCGA-OV **(f)** cohorts, stratified into high (red) and low (blue) expression groups based on the median score of the CAF-like signature.
**g, h,** Kaplan–Meier overall survival curves for TCGA-STAD patients stratified into high (red) and low (blue) expression groups based on the median score of the CAF-like signature across tumor samples **(g)**, and adjacent normal tissue samples **(h)**.
**i,** Forest plots of univariate Cox proportional hazards regression analysis for individual CAF-like marker genes in TCGA-STAD tumor and adjacent normal tissue. Hazard ratios (HRs) are presented per standard deviation increase in gene expression. Points represent HR estimates, and horizontal bars indicate 95% confidence intervals. Red markers indicate statistically significant associations ($P < 0.05$), whereas blue markers indicate non-significant associations.
